# Identification and characterization of small molecule inhibitors of the LINE-1 retrotransposon endonuclease

**DOI:** 10.1101/2022.12.29.522256

**Authors:** Alexandra M. D’Ordine, Gerwald Jogl, John M. Sedivy

**Author notes:** Correspondence: John Sedivy,; Gerwald Jogl.

## Abstract

The long interspersed nuclear element-1 (LINE-1 or L1) retrotransposon is the only active autonomously replicating retrotransposon in the human genome. While most of the approximately 500,000 L1 copies are no longer able to propagate, those that retain activity can harm the cell by creating mutations, generating DNA damage, and triggering the expression of inflammatory factors such as the host interferon anti-viral response. Therefore, inhibition of L1 could be used to treat a variety of diseases associated with these processes. Previous research has focused on inhibition of the L1 reverse transcriptase (RT) activity, in part due to the prevalence of well-characterized existing inhibitors to related viral enzymes. Here we present the L1 endonuclease (EN) as an additional target for reducing the detrimental effects of L1 expression. We have screened and characterized a set of structurally diverse small molecule EN inhibitors using computational, biochemical, and cellular methods. We also show that these inhibitors reduce DNA damage created by L1 and inflammation reinforced by L1 activity in senescent cells. These inhibitors could be further used to modulate endogenous L1 function to better understand the lifecycle of this ubiquitous disease-relevant element.

## Introduction

The long interspersed nuclear element-1 (LINE-1 or L1) retrotransposon comprises approximately 17% of the human genome^1^. This sequence propagates using an RNA intermediate in a “copy and paste” mechanism known as retrotransposition. L1 is the only active autonomously replicating retrotransposon in humans, encoding the proteins required for its retrotransposition^2^. While most L1 sequences in the human genome are truncated or mutated, a small number (80-100) retain the ability to create additional L1 insertions^3^ and mobilize other non-autonomous retrotransposons, Alu and SVA, which make up another 11% of the human genome^1^. Activity of these elements is associated with genomic instability since they can create deletions, inversions, and other rearrangements, in addition to mutations and insertions resulting from integration of new copies^4,5^. New L1 or L1-driven Alu/SVA germline insertions occur in up to one in every 20 individuals^6^ and can cause disease^7^. Somatic L1 activity may function during development^8,9^, but has also been associated with cancer and neurodegeneration^10,11^. Most recently, L1 expression has been shown to reinforce the inflammatory phenotype of senescent cells^12^, which are cells that have permanently exited the cell cycle and contribute to sterile inflammation during aging.

L1 encodes three proteins: ORF0, ORF1, and ORF2. ORF0 is a 7 kDa primate-specific protein of unknown function located on the antisense strand^13^. ORF1 is a 40 kDa trimeric RNA-binding protein that binds the L1 transcript and is required for L1 retrotransposition^14,15^. ORF2 is a 150 kDa multi-functional protein that contains the enzymatic activities needed for retrotransposition: an N-terminal apurinic/apyrimidinic (AP)-like endonuclease (EN) domain^16^, a reverse transcriptase (RT) domain^17^, and a C-terminal cysteine-rich region of unknown function^18^. ORF1 and ORF2 preferentially bind the L1 transcript in *cis* to form a ribonucleoprotein (RNP) particle^19^. The L1 RNP particles contain many copies of ORF1 and only one or two copies of ORF2, but the stoichiometry needed for retrotransposition is unknown^20^. To insert a new copy into the genome, L1 uses target-primed reverse transcription (TPRT)^2^. The EN initiates this pathway by creating a single-stranded nick in genomic DNA at the semi-specific 5′-TTTT*A-3′ consensus sequence^21^, with the asterisk marking the location of the cleaved phosphodiester bond. The exposed poly-T sequence base-pairs with the poly-A tail of the L1 transcript to allow for priming of reverse transcription by the RT to create L1 complementary DNA (cDNA). This step is followed by a second nick, possibly created by the EN, and polymerization of the second L1 strand. Finally, host factors integrate the new copy of L1, resulting in flanking target site duplications, a hallmark of canonical TPRT. Both the EN and RT are required for retrotransposition, as active site mutations abolish retrotranposition^14,16^, though EN-independent retrotransposition can occur in cells deficient for non-homologous end joining and at dysfunctional telomeres^22,23^. Therefore, the EN plays a key role in beginning the process of creating and integrating a new L1 copy into the genome.

Previous studies have shown that pharmacological inhibition of the RT reduces retrotransposition to a similar extent as active site mutation^24,25^. These inhibitors also decrease the amount of pro-inflammatory L1 cDNA found in the cytoplasm of senescent cells^12,26^. However, no small molecule EN inhibitors have been characterized. The EN is a promising target for several reasons. The crystal structures of the EN alone^27^ and bound to substrate DNA^28^ have been solved, enabling use of *in silico* screening of candidate small molecules. A similar approach was used to successfully isolate inhibitors of APE-1^29,30^, a structurally related enzyme^27^. The EN is also amenable to biochemical characterization and *in vitro* inhibitor screening. Additionally, the DNA damage and cytotoxicity induced by L1 activity is driven in part by the EN. Expression of L1 causes DNA damage demonstrated by accumulation of γ-H2AX^31–34^ and 53BP1 foci^33,35^, as well as overall DNA fragmentation by neutral COMET assay^31,32,35^. Significantly, expression of ORF2 without ORF1^32,34,35^, or of the EN alone ^32,36^, results in DNA damage. This DNA damage is impaired in the case of EN mutation^31,32,36^ and even more so when both the RT and EN are mutated^32–34^. Expression of ectopic L1 elements can reduce cell viability, which is partially rescued when EN is mutated, in some cases to a larger extent than RT mutation alone^31,37^. This cytotoxicity also occurs with expression of ORF2 or the EN alone^32,36,37^. These results demonstrate that not only does L1 create DNA damage and cytotoxicity, but that both can occur in the absence of actual retrotransposition and appear to be largely due to EN activity. In fact, it has been suggested that up to 10 times more double-strand breaks occur in cells overexpressing L1 than productive insertions based on relative frequencies of γ-H2AX foci and retrotransposition events^31^. Furthermore, EN inhibitors would be very useful to understand EN function in the context of natural L1 lifecycles, since our current knowledge is largely based on studies using ectopically introduced L1 overexpression constructs. Specifically, selective inhibition of the EN in senescent cells could help elucidate the mechanism of cytoplasmic L1 cDNA formation and subsequent triggering of the type I interferon (IFN-I) response. Finally, development of EN inhibitors would enable combining pharmacological inhibition of both catalytic L1 domains for potential synergistic effects.

## Results

### Identification of initial EN inhibitors

In the absence of any existing EN small molecule inhibitors, we first explored inhibitors of the structurally related human endonuclease APE-1^29^, which functions in base excision repair. This approach was similarly used to discover that some HIV RT inhibitors, such as 3TC, also have efficacy against the L1 RT and can be repurposed for L1 inhibition^24,25^. Using the molecular docking program LeDock^38^ and the crystal structure of the L1 EN^27^ (PDB ID: 1VYB), we evaluated 15 reported APE-1 inhibitors. We chose to test one of the weaker APE-1 inhibitors (K_i_ for APE-1=13μM^29^), NSC89640, as it resulted in the most favorable docking score (−8.39 kJ/mol). We also obtained the APE-1 inhibitor with the best efficacy against APE-1, NSC332395 (K_i_ for APE-1=0.12μM^29^), though it was predicted to have a less favorable docking score for EN (−5.75 kJ/mol). We evaluated these two preliminary candidate compounds using the established plasmid nicking assay for L1 EN activity^16^. In this assay, activity is visualized as the slower migration of nicked versus supercoiled plasmids by agarose gel electrophoresis. We found that NSC332395 did not inhibit EN activity in this assay, whereas NSC89640 (referred to now as AD2), resulted in full inhibition at 1mM (Fig 1A, Table 1). While this constituted weak inhibition according to this qualitative activity assay, we used AD2 as a preliminary inhibitor for finding additional compounds for testing.

**Table 1:**
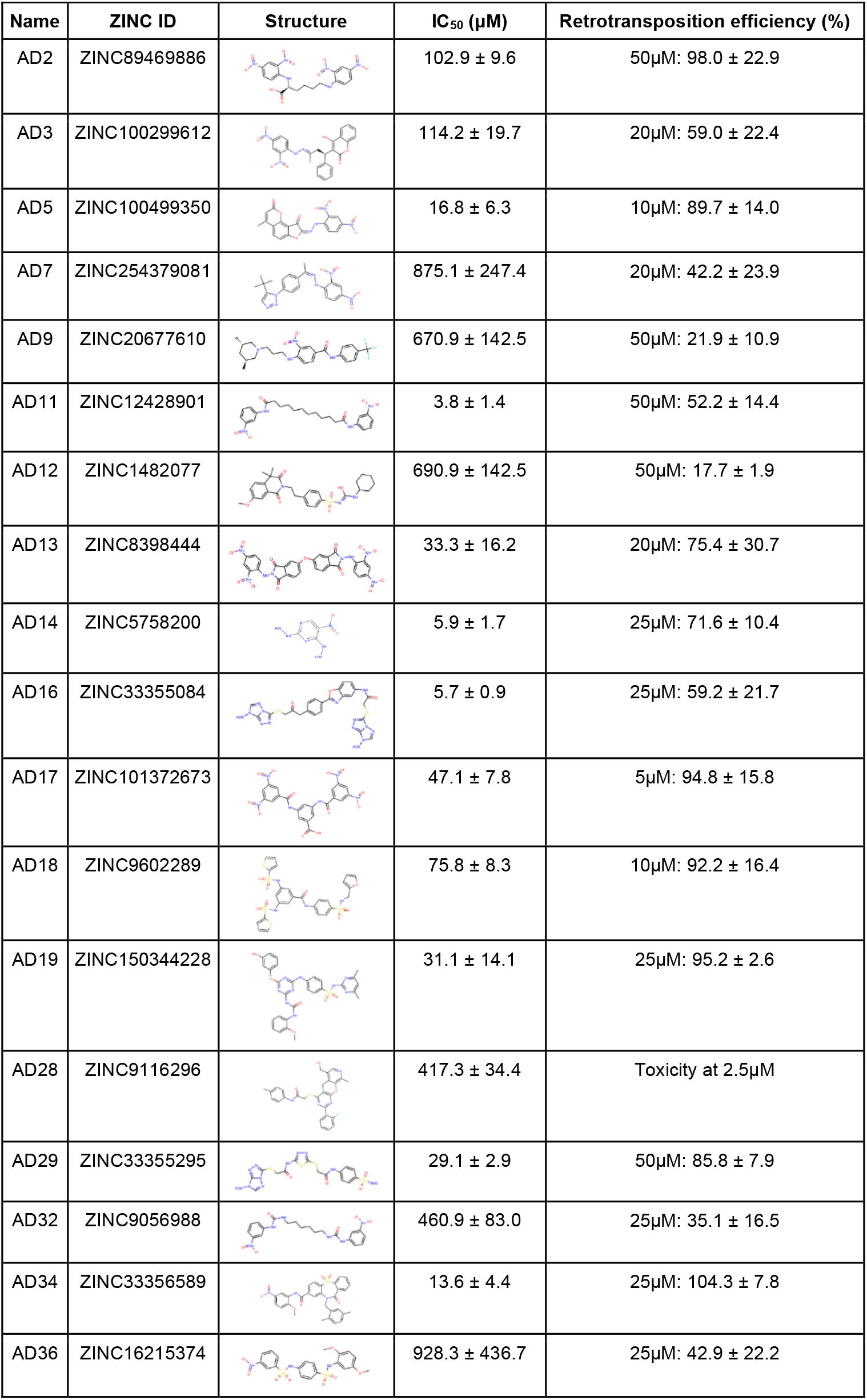
Summary of EN inhibitors. Structure images from the ZINC database. IC_50_ values were calculated from 3 independent experiments of the fluorescent oligonucleotide nicking assay with 3 replicates each and are mean ± S.D. Retrotransposition efficiencies relative to no inhibitor control were calculated from at least 3 independent experiments with 4 replicates each and are mean ± S.D.

| Name | ZINC ID | Structure | IC<sub>50</sub> (μM) | Retrotransposition efficiency (%) |
| --- | --- | --- | --- | --- |
| AD2 | ZINC89469886 | | 102.9 ± 9.6 | 50μM: 98.0 ± 22.9 |
| AD3 | ZINC100299612 | | 114.2 ± 19.7 | 20μM: 59.0 ± 22.4 |
| AD5 | ZINC100499350 | | 16.8 ± 6.3 | 10μM: 89.7 ± 14.0 |
| AD7 | ZINC254379081 | | 875.1 ± 247.4 | 20μM: 42.2 ± 23.9 |
| AD9 | ZINC20677610 | | 670.9 ± 142.5 | 50μM: 21.9 ± 10.9 |
| AD11 | ZINC12428901 | | 3.8 ± 1.4 | 50μM: 52.2 ± 14.4 |
| AD12 | ZINC1482077 | | 690.9 ± 142.5 | 50μM: 17.7 ± 1.9 |
| AD13 | ZINC8398444 | | 33.3 ± 16.2 | 20μM: 75.4 ± 30.7 |
| AD14 | ZINC5758200 | | 5.9 ± 1.7 | 25μM: 71.6 ± 10.4 |
| AD16 | ZINC33355084 | | 5.7 ± 0.9 | 25μM: 59.2 ± 21.7 |
| AD17 | ZINC101372673 | | 47.1 ± 7.8 | 5μM: 94.8 ± 15.8 |
| AD18 | ZINC9602289 | | 75.8 ± 8.3 | 10μM: 92.2 ± 16.4 |
| AD19 | ZINC150344228 | | 31.1 ± 14.1 | 25μM: 95.2 ± 2.6 |
| AD28 | ZINC9116296 | | 417.3 ± 34.4 | Toxicity at 2.5μM |
| AD29 | ZINC33355295 | | 29.1 ± 2.9 | 50μM: 85.8 ± 7.9 |
| AD32 | ZINC9056988 | | 460.9 ± 83.0 | 25μM: 35.1 ± 16.5 |
| AD34 | ZINC33356589 | | 13.6 ± 4.4 | 25μM: 104.3 ± 7.8 |
| AD36 | ZINC16215374 | | 928.3 ± 436.7 | 25μM: 42.9 ± 22.2 |

**Figure 1:**
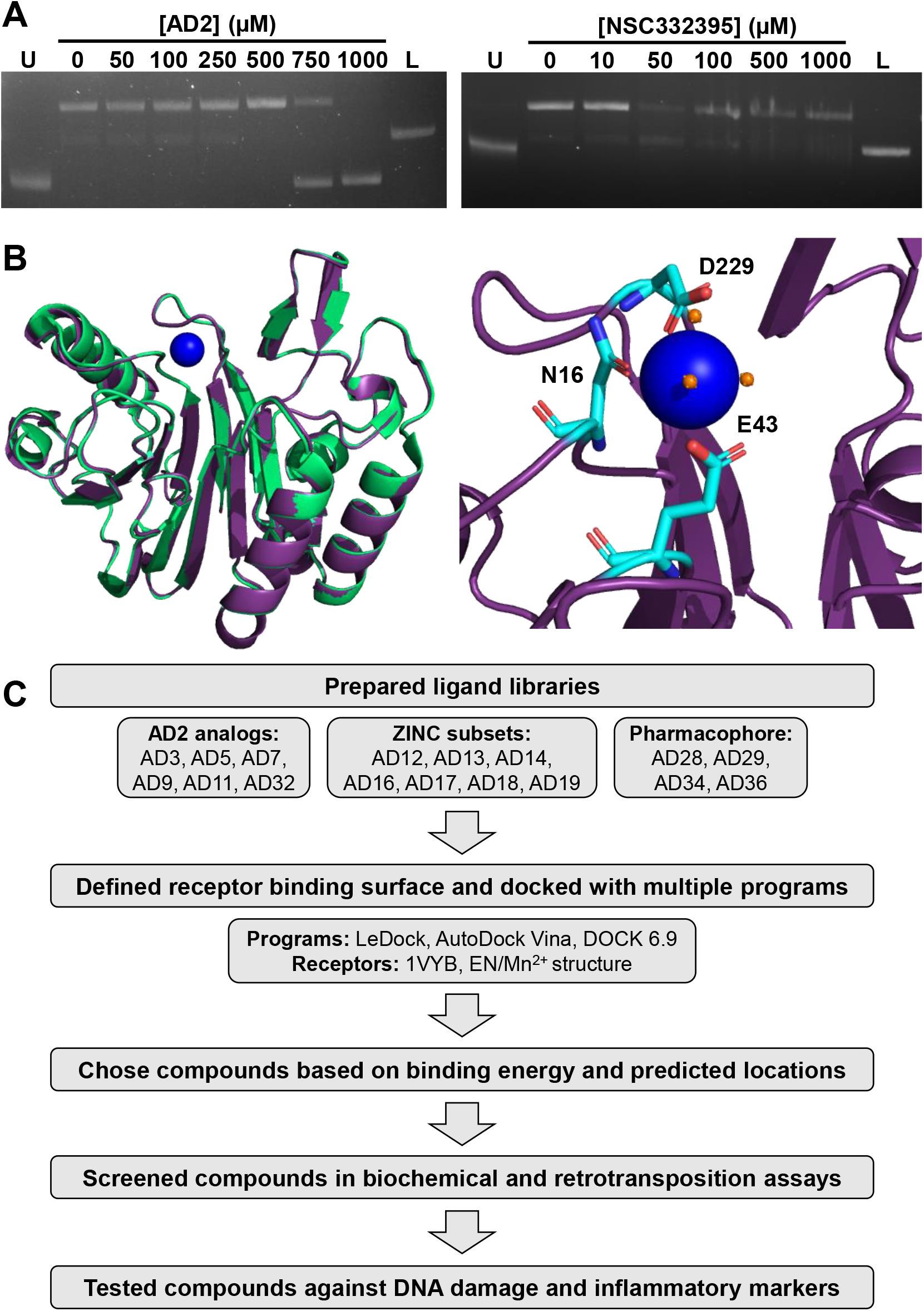
**A)** Plasmids nicking assay results for AD2 (left) and NSC332395 (right). U; uncut plasmid, L; linearized plasmid. **B)** EN crystal structure with Mn^2+^ ion bound. Left: Overlay of new structure (EN, purple; Mn^2+^, blue) with published EN structure (green, PDB ID: 1VYB), RMSD= 0.254. Right: Coordination of Mn^2+^ with indicated active site residues (light blue) and waters (orange). **C)** Summary of compound screening strategy and origins of EN inhibitors.

### Evaluating small molecules *in silico*

We performed multiple rounds of screening involving docking to find EN inhibitors: structural analogs of AD2 and subsequent inhibitors, additional subsets from the ZINC database, and compounds from pharmacophore filtering. Using the ZINC database^39^, we selected structural analogs of AD2 and evaluated them using LeDock^38^, AutoDock Vina^40^, and DOCK 6.9^41^. By using multiple algorithms to screen compounds, we aimed to increase the accuracy of our predictions by choosing compounds with favorable binding energies and similar binding positions in the active site. Next, we decided to expand the structural diversity of compounds screened. For this screening we solved a crystal structure of the EN bound to a Mn^2+^ ion (Fig. 1B) and used this as the receptor to better represent the electrostatic environment of the active site. This new structure enabled us to find additional inhibitors using LeDock and AutoDock Vina by screening ZINC subsets, including FDA and internationally approved drugs; previous attempts using the apo EN structure to dock compound libraries unbiased by AD2 resulted in no compounds with *in vitro* efficacy. Finally, we generated a structure-based pharmacophore using a structure of the EN bound to substrate dsDNA^28^ (PDB ID: 7N94). We used ZINCPharmer^42^ to generate the pharmacophore and filter the ZINC database for compounds that matched the pharmacophore, followed by docking with LeDock and AutoDock Vina as previously described. A summary of docking and the subsequent inhibitor testing workflow is shown in Fig. 1C.

### Fluorescent oligonucleotide biochemical assay to quantify efficacy of EN inhibitors

We used two assays to biochemically characterize candidate EN inhibitors. We initially tested 15 AD2 analogs with the plasmid nicking assay, and found several compounds with inhibition (Fig. 2A, Table 1). While this assay provides qualitative evidence for inhibition, we sought a more quantitative assay in order to prioritize inhibitors for future development and testing in cells. We adapted a fluorescent hairpin oligonucleotide assay used for APE-1^29^ for use with EN (Fig. 2B). We replaced the abasic site analog and surrounding sequence with the EN target sequence 5’-TTTTA-3’. EN activity releases the 5’ 8-nucleotide fragment with a fluorescent tag that dissociates from the remaining sequence containing the 3’ quencher. This allows for real-time monitoring of activity based on fluorescence intensity and quantification of initial reaction rates across multiple inhibitor concentrations. We then utilized this assay to screen subsequent rounds of inhibitors and to calculate IC_50_ values (Fig. 2C, Table 1). These results show EN inhibitors with diverse structural scaffolds with potency down to the low micromolar range.

**Figure 2:**
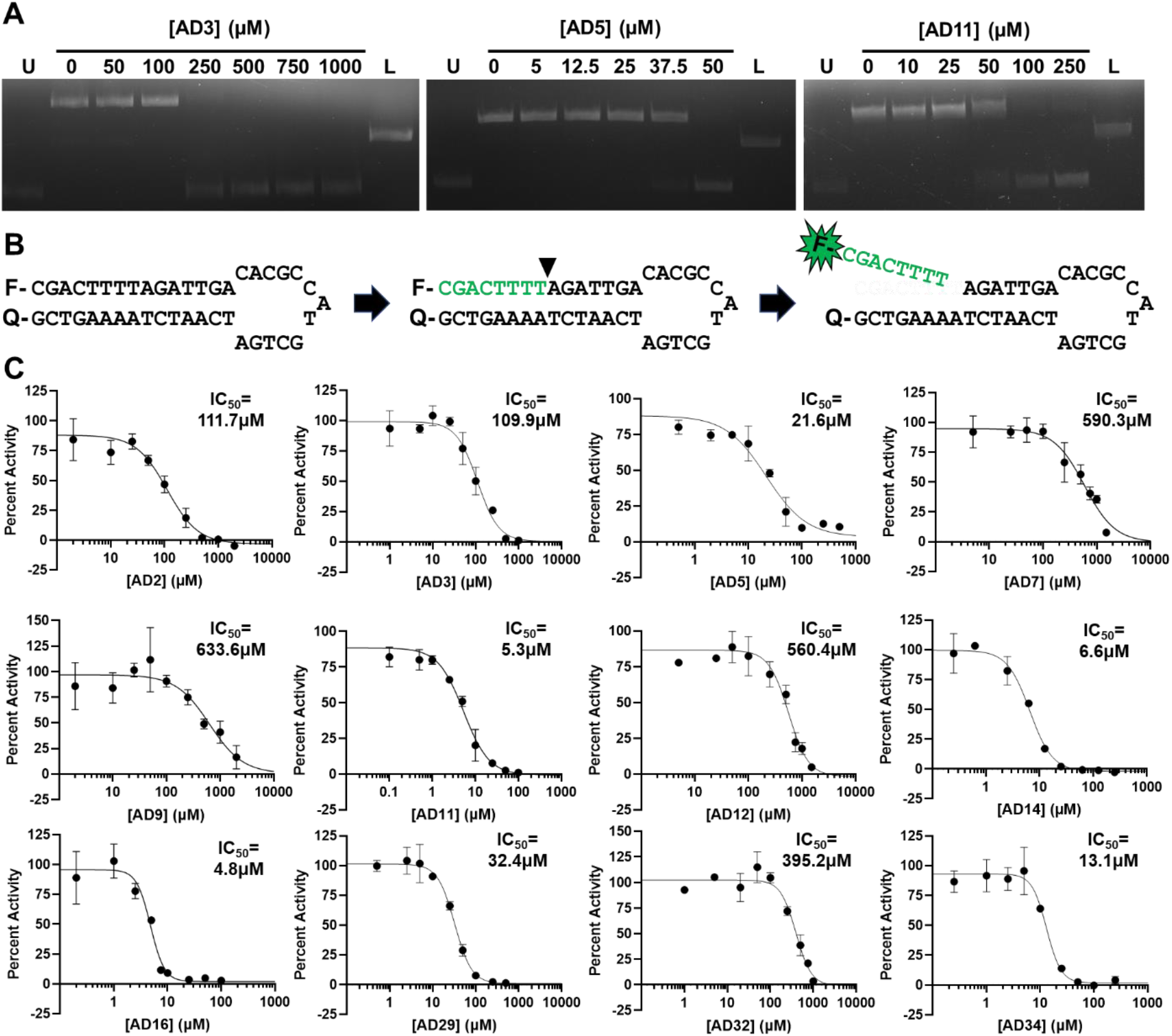
Biochemical characterization of EN inhibitors. **A)** Plasmid nicking assay results for selected AD2 analogs. U; uncut plasmid, L; linearized plasmid. **B)** Schematic of fluorescent oligonucleotide nicking assay. Left: sequence of hairpin oligonucleotide containing 5’ 6-FAM fluorescein fluorophore (F) and 3’ DABCYL quencher (Q). Middle: Arrowhead indicates location of nick by EN at the semi-specific target site sequence 5’-TTTT*A-3’. Melting temperature of green sequence is lower than reaction temperature, whereas melting temperature of full hairpin is higher than the reaction temperature. Right: Nicked sequence is released away from the quencher, allowing for fluorescence to occur as a real-time readout for activity using a plate reader. **C)** Representative assay results for EN inhibitors. Graphs show percent of no inhibitor control EN activity as a function of indicated inhibitor concentration. Activity was determined as initial rate of reaction under multiple turnover conditions and normalized to no inhibitor control. IC_50_ values were calculated using [inhibitor] vs. response non-linear fit in GraphPad Prism version 9.4.1 for Windows. No inhibitor control and full inhibition by 50mM EDTA were included in fit calculations to guide definition of top and bottom of fit curve.

### EN inhibitors reduce L1 retrotransposition in cell culture

In addition to measuring EN inhibition *in vitro*, we determined the effects of these inhibitors on L1 retrotransposition in cells. We used an established HeLa cell culture dual-luciferase reporter assay to measure L1 retrotransposition^14,43^. In this assay, expression of a luciferase reporter occurs only after insertion of a new L1 element. Several compounds that inhibited EN activity in the *in vitro* assays reduced retrotransposition relative to the no inhibitor control (Fig. 3A). The RT inhibitor 3TC served as a comparison to inhibition of the RT domain. Several EN inhibitors decreased retrotransposition to a similar degree as 3TC. Inhibition using multiple concentrations of selected inhibitors is shown in (Fig. 3B). A summary of EN inhibitor structures, IC_50_ values, and retrotransposition efficiencies across multiple independent experiments is shown in Table 1.

**Figure 3:**
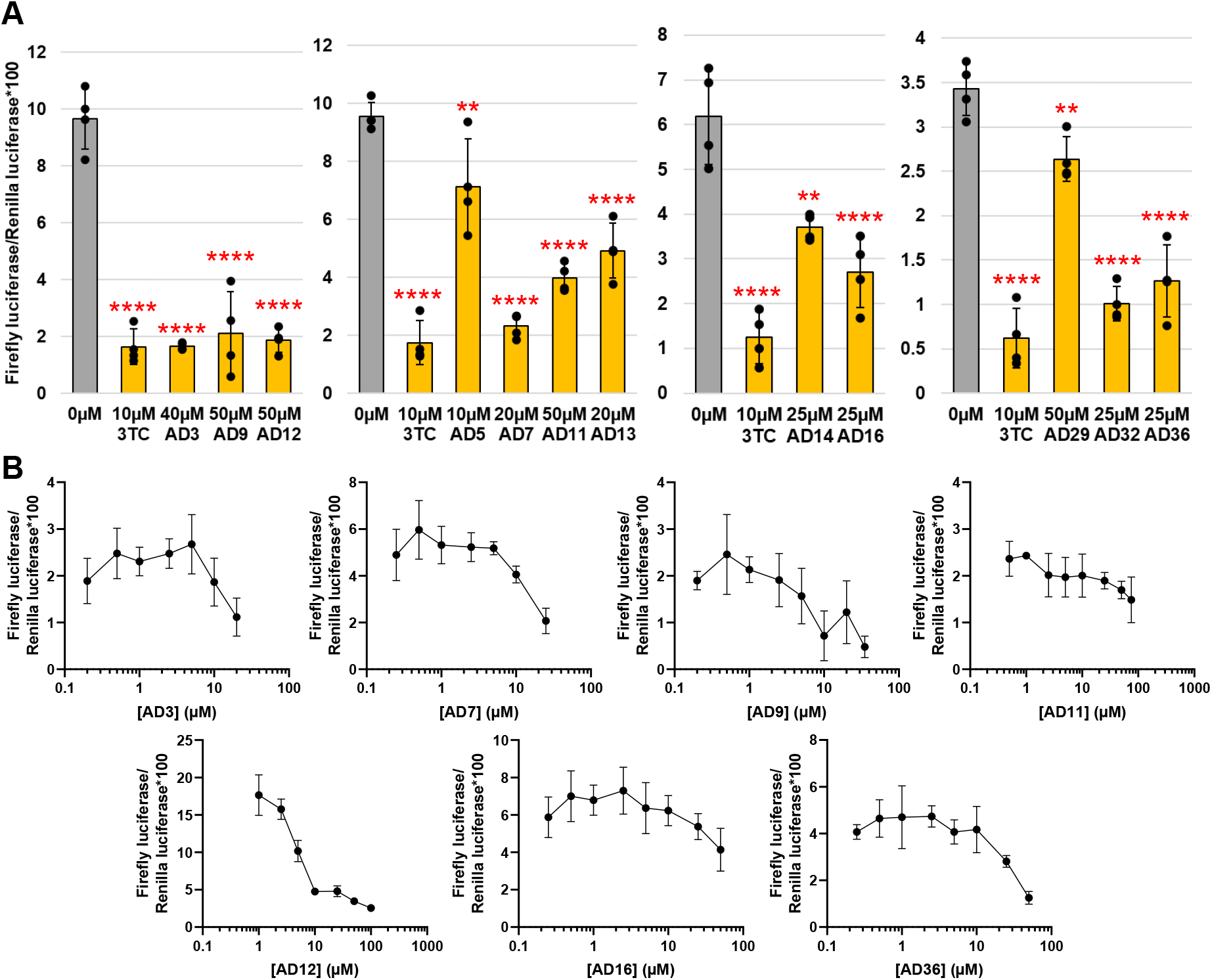
Effects of EN inhibitors on L1 retrotransposition in cell culture. **A)** Retrotransposition was measured in HeLa cells by expression of a Firefly luciferase reporter after a retrotransposition event normalized to Renilla luciferase activity. 3TC was included for comparison to RT inhibition. During compound screening, PrestoBlue Viability Reagent was used to test cytotoxicity; compounds with statistically significant cytotoxicity were subsequently tested at lower concentrations or excluded from further testing. Statistical significance of the mean relative to no inhibitor control (0µM) was determined by one-way ANOVA followed by Dunnett’s multiple comparisons test using GraphPad Prism version 9.4.1 for Windows. *p<0.05, **p<0.01, ***p<0.001, ****p<0.0001. **B)** Concentration-dependent inhibition for selected EN inhibitors. Results from 4 replicates for each treatment and concentration. Error bars=S.D.

### L1-induced DNA damage is mitigated by EN inhibitors

After determining the effects of EN inhibitors on L1 retrotransposition, we examined other phenotypes resulting from L1 expression. Inactivation of EN by active site point mutation has been previously shown to significantly reduce DNA damage when L1 is overexpressed^31,32,36^. We thus transfected inducible plasmid constructs expressing full-length L1 (FL, ORF1 and ORF2) or the EN domain only into HeLa Tet-On cells. After induction of expression with doxycycline and addition of inhibitors, DNA damage was detected using immunofluorescence for γ-H2AX, a marker of double-stranded DNA breaks (Fig. 4A). This staining was then quantified using a pipeline from CellProfiler that we adapted to measure the average γ-H2AX signal for each nucleus identified. Our results show that the inhibitors reduce γ-H2AX signal both when FL L1 is expressed, as well as when the EN domain is expressed alone (Fig. 4B). We have also qualitatively shown by neutral COMET assay that EN inhibitors can mitigate overall DNA fragmentation caused by EN domain expression (Fig. 4C). Our results demonstrate that small molecule EN inhibitors can reduce DNA damage caused by the expression of L1 elements.

**Figure 4:**
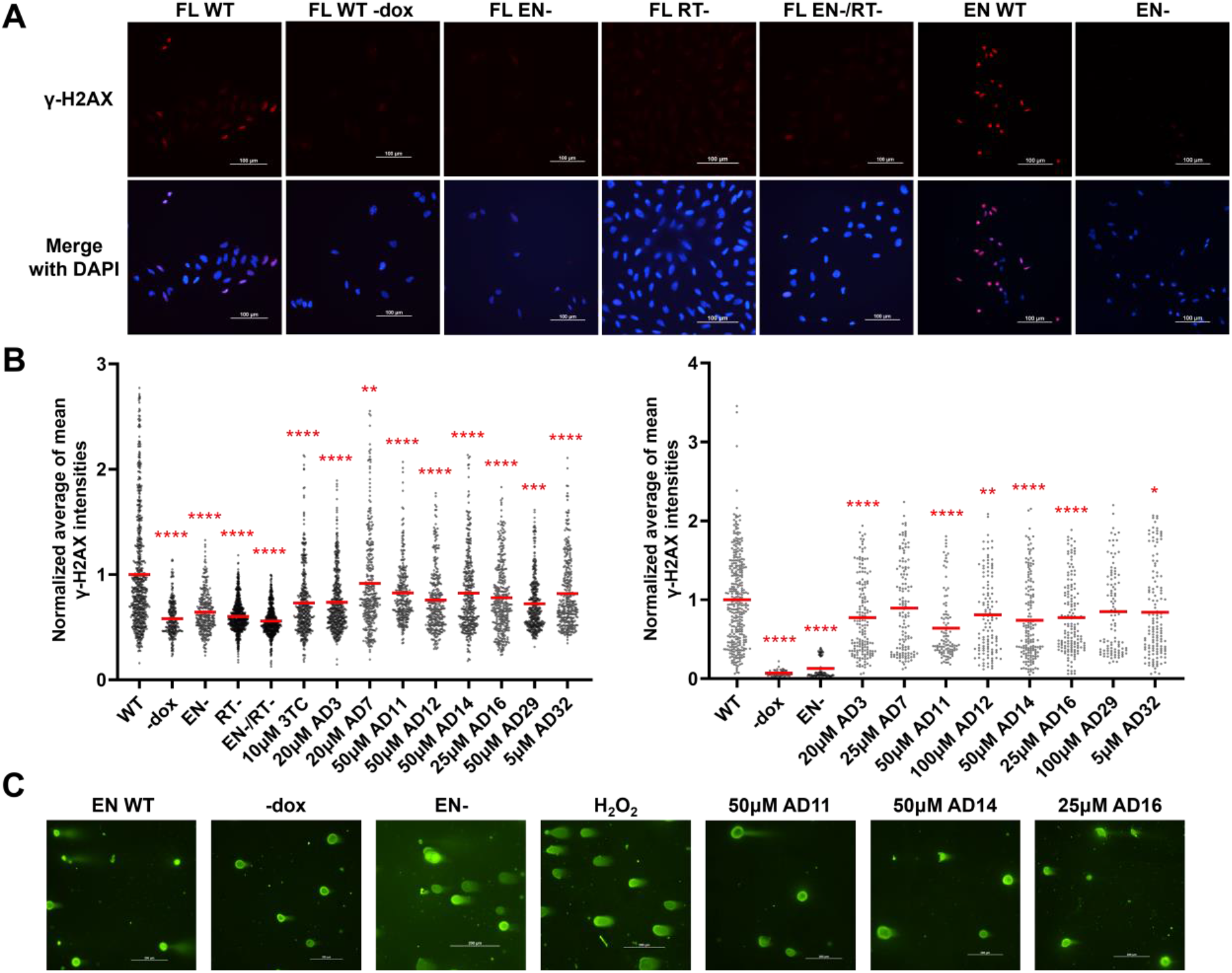
Effects of EN inhibitors on L1-induced DNA damage. HeLa Tet-On cell lines containing doxycycline-inducible FL or EN domain only L1 expression constructs were generated for WT and mutant (EN-, H230A; RT-, D702Y) L1. **A)** Representative immunofluorescence images of HeLa cells with indicated constructs stained for γ-H2AX. **B)** Average of mean γ-H2AX intensities of individual nuclei normalized to WT no inhibitor control for FL expression (left, n=254-752) or EN domain (right, n=56-325). Results are from at least 3 independent experiments per sample. All samples from the same experiment were processed in parallel and images were acquired with the same exposure. Mean γ-H2AX intensity analysis was performed with CellProfiler. Statistical significance of mean (red bars) vs. WT was determined by one-way ANOVA followed by Dunnett’s multiple comparisons test using GraphPad Prism version 9.4.1 for Windows. *p<0.05, **p<0.01, ***p<0.001, ****p<0.0001. **C)** Representative images of neutral COMET assay results for HeLa cells expressing EN domain constructs and treated with inhibitors or hydrogen peroxide as indicated.

### EN inhibitors impact senescence-associated inflammatory markers in cell culture

We tested the effects of EN inhibitors on the expression of inflammatory markers in senescent cells. To perform these experiments, we used a replicative senescence cell culture model (Fig. 5A). Human diploid fibroblasts were passaged until they no longer were dividing, then maintained in culture for at least 4 months. This duration was based on previous evidence from our laboratory demonstrating significant expression of L1 at later stages of senescence beginning around 3 months^12^. We then treated the cultures with inhibitors for 1 month before measuring levels of inflammatory markers by RT-qPCR. We found similar effects for 3TC and EN inhibitors on the indicated markers in two independent cultures of senescent cells (Fig. 5B, Fig. 5C).

**Figure 5:**
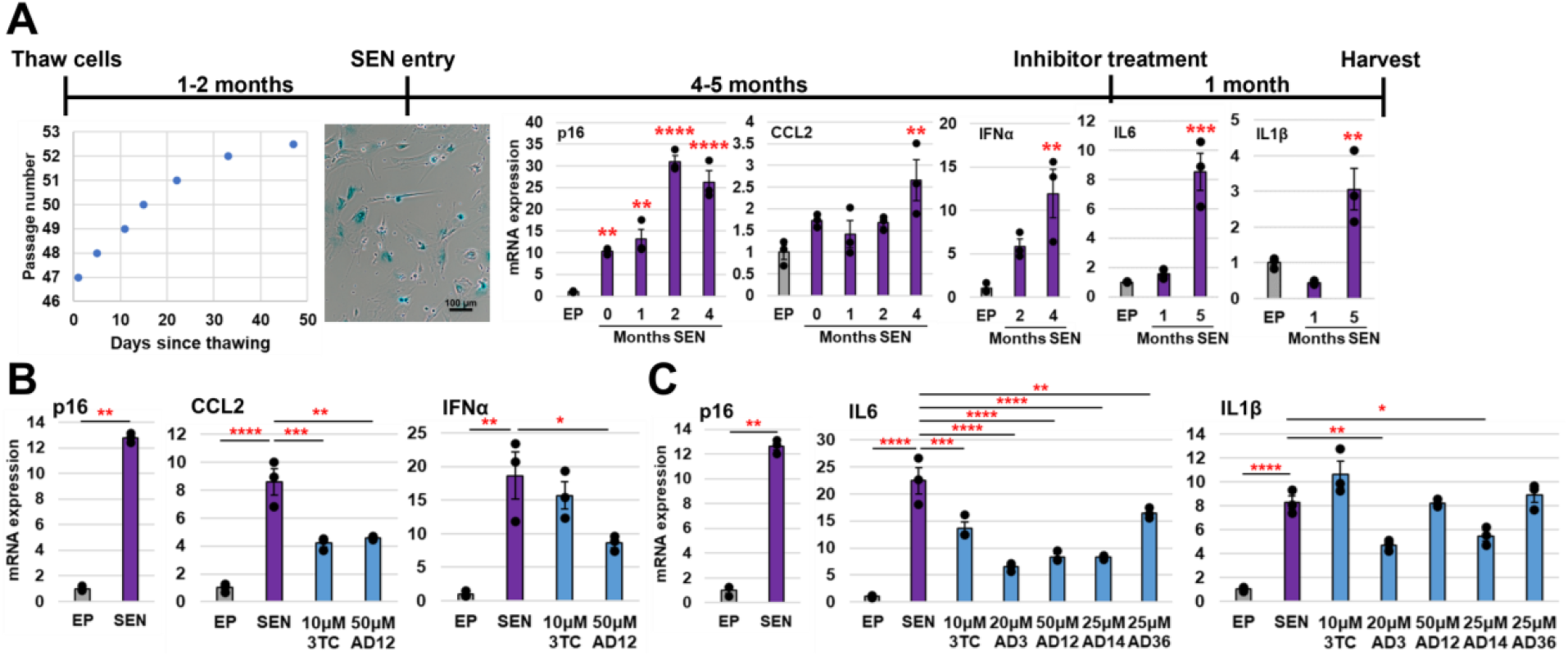
EN inhibitors impact inflammation markers in senescent cells. **A)** Timeline of senescent (SEN) culture generation and treatment. Cells were passaged until they stop dividing and then maintained for 4 to 5 months, after which they were treated with inhibitors for 1 month before harvesting for RNA. Left: Representative passaging history showing progressive slowing of cell division. Middle: Senescence-associated β-galactosidase assay of cells at SEN entry (0 months SEN) with blue staining indicating senescence. Right: Representative RT-qPCR results of p16, SASP markers IL6, IL1β, CCL2, and MMP3, and IFN-I marker IFNα throughout senescence normalized to early passage (EP) proliferating cells. **B)** RT-qPCR of SEN culture treated with inhibitors from 4 to 5 months SEN normalized to EP. **C)** RT-qPCR of SASP markers IL6 and IL1β for SEN culture treated with inhibitors from 5 to 6 months SEN normalized to EP. Statistical significance of the mean relative to EP (A) or SEN (B,C) was determined by one-way ANOVA followed by Dunnett’s multiple comparisons test using GraphPad Prism version 9.4.1 for Windows. Statistical significance for p16 in (B,C) was determined using an unpaired two-tailed *t*-test. Error bars=S.E., *p<0.05, **p<0.01, ***p<0.001, ****p<0.0001.

## Discussion

We describe here a set of structurally diverse small molecule L1 EN inhibitors. We have identified these inhibitors through various computational screening methods and quantified their respective efficacies biochemically as well as in cells. Furthermore, we have demonstrated that these inhibitors mitigate L1-induced DNA damage and inflammation in senescent cells, which are disease-relevant impacts of L1 activity. These inhibitors could serve as key tools in better understanding L1 function from a basic science perspective and as initial candidates for the development of therapeutics for age-associated diseases.

We found clear differential potency among the inhibitors when assessed by the various downstream assays. Several inhibitors showed clear inhibition in the fluorescent oligonucleotide nicking assay but did not significantly impact retrotransposition in cells: AD2, AD5, AD17, AD18, AD19, AD34. This discrepancy could be due to several factors including low solubility or poor cell permeability. Among these inhibitors, AD18 and AD19 could be good starting points for further development of compounds with improved solubility and cell permeability, since they already have good *in vitro* potency and have unique functional groups relative to the other inhibitors. On the other hand, several inhibitors showed relatively low efficacy *in vitro* but high activity in the retrotransposition assay: AD7, AD9, AD12, AD32, and AD36. One possibility is that these compounds have higher cell permeability, allowing them to achieve higher intracellular concentrations. This is supported by the fact that AD12 is a therapeutic for diabetes mellitus type 2 (Gliquidone) and therefore has been optimized for bioavailability. In a biological context, even weak EN inhibition might significantly impact retrotransposition. First, TPRT only requires one or two nicks by the EN for productive insertion, a situation clearly different from the multi-turnover kinetics we measured *in vitro*. Second, the EN is a relatively slow enzyme^16^, similar to retroviral integrases^44^ that also only need to accomplish a few catalytic cycles for propagation. This might be an evolutionary advantage that allows for a balance between L1 integration and EN-dependent DNA damage that could harm the host. Finally, it has to be considered that *in vivo* EN activity occurs within a complex L1 RNP that interacts with multiple host proteins in the nucleus^20,45^. Some evidence suggests that EN activity is reduced in full length ORF2^46^, potentially resulting in partial occlusion of the DNA and therefore inhibitor binding surface. Conversely, the full length ORF2 could provide additional binding surfaces for inhibitor interactions that could reduce retrotransposition by impairing ORF2 function overall, including the RT domain directly or the coordination between the domains during TPRT.

The inhibitors also impacted L1 activity differently in the various cell-based assays used here. While the retrotransposition assay requires full length L1, the DNA damage assays were performed using both full length L1 and EN domain-only constructs. This allows for evaluation of inhibition of DNA damage in the content of L1 retrotransposition-competent full length sequences in comparison to the EN domain alone. L1 elements that are incapable of retrotransposition but retain potentially active EN coding sequences are abundant in the human genome and thus have the potential to create DNA damage if expressed^32^. It is interesting to note that inhibitors with the best inhibition in the full length L1 DNA damage experiments were generally the ones with the best inhibition in the retrotransposition assay, for example AD7 and AD12. In a similar way, inhibitors with better efficacy in the EN domain-only DNA damage experiments were more likely to have lower *in vitro* IC50 values, such as AD11 and AD14. This suggests that various inhibitors have the potential to interact differently in the context of the full length ORF2 versus the EN domain alone.

Our results regarding the role of the EN in retrotransposition and DNA damage agree with previous research demonstrating that when overexpressing L1, EN active site mutations prevent retrotransposition and DNA damage^16,31,32,36,47,48^. However, analogous experiments have not been performed to investigate the role of the EN in the production of inflammatory L1 cDNA found in the cytoplasm of senescent cells. This is mostly because of technical challenges inherent in creating active site mutations in endogenous L1s being expressed from multiple loci in the genome. RT inhibitors like 3TC have helped answer these questions with regard to RT function^12^, but the absence of EN inhibitors has prevented similar experiments to investigate the role of the EN. We report here early data on testing the role of EN in the pathways that generate these inflammatory signals. Our results suggest that the EN is at least partially involved in this process, as EN and RT inhibition resulted in similar decreases in expression of some SASP markers. One possibility is that nuclear envelope damage that has been widely documented in senescent cells^49,50^ allows for intermediates of retrotransposition primed in the nucleus under the conditions of canonical TPRT to enter the cytoplasm and trigger the IFN-I response through the cGAS-STING pathway^51^. Another possible explanation is that the EN can function in the cytoplasm to improve RT priming by nicking chromatin fragments prevalent in senescent cells^52,53^ or other sequences such as mitochondrial DNA, with preference for AT-rich sequences that promote RT priming^54^. However, these hypotheses require further testing using a multitude of approaches.

We believe the most promising inhibitors for further exploration based on structural diversity and efficacy across assays to be AD3, AD11, AD12, AD14, and AD16. Broadly speaking, these inhibitors could be used to study EN function in a wider range of *in vivo* contexts than currently possible. As previously noted, most of what we know about L1 has been garnered using overexpression models. While useful, including for experiments in this study, this approach should be complemented with more biologically relevant ones that rely on endogenous L1 expression. These inhibitors could therefore be used to determine the effects of EN inhibition across cell lines, animal models, and diseases, such as in cancer and neurodegeneration where retrotransposition, DNA damage, and inflammation resulting in part from L1 may all play roles in disease progression. EN inhibitors specifically provide an orthogonal pharmacological approach to L1 inhibition. RT inhibitors, while available and numerous, mostly rely on the same chain-terminating chemistry and therefore have similar off-target effects. The variety of structures of EN inhibitors described here, in combination with the relatively large DNA-binding surface of the EN, suggests that more classes of EN inhibitors are theoretically possible. This could mitigate concerns and provide better management of off-target effects. Finally, EN inhibitors could be advantageous for reducing DNA damage specifically, as this domain is directly responsible for DNA damage by nicking DNA, which in some contexts could be independent of insertion events.

In summary, we have characterized the first set of small molecule inhibitors of the LINE-1 retrotransposon endonuclease domain. In the short term, these inhibitors can serve as tools to improve our understanding of L1 biology in a similar way to how RT inhibitors have been used. Ultimately, these inhibitors represent a starting point for future development of potential therapeutics for diseases associated with L1 expression, many of which are likely to be age-associated diseases.

## Methods

### Materials

NSC332395 was obtained from the National Cancer Institute. The following EN inhibitor was obtained from Sigma: AD12/ZINC1482077 (CDS021537). The following EN inhibitors were obtained from the listed suppliers through Molport: AD2/ZINC89469886 (Alinda IBS-L0127235), AD3/ZINC100299612 (ChemBridge 5151622), AD5/ZINC100499350 (ChemDiv 1440-2881), AD7/ZINC254379081 (Enamine Z56821059), AD9/ZINC20677610 (ChemDiv K784-1448), AD11/ZINC12428901 (ChemDiv 8003-9274), AD13/ZINC8398444 (Specs AG-690/11231133), AD14/ZINC5758200 (Specs AC-907/25004307), AD16/ ZINC33355084 (Vitas STK672667), AD17/ZINC101372673 (Vitas STK000838), AD18/ZINC96022289 (UkrOrgSynthesis PB56889488), AD19/ZINC150344228 (TimTec ST026710), AD28/ZINC9116296 (ChemDiv E544-0411), AD29/ZINC33355295 (ChemDiv 8008-0573), AD32/ZINC9056988 (TimTec ST002110), AD34/ZINC33356589 (Specs AQ-088/42014071), AD36/ZINC16215374 (ChemBridge 7937857). HeLa Tet-On cells containing plasmid pPM404^55^, which expresses the L1 WT *ORFeus* sequence with a dual-luciferase reporter for use in the L1 retrotransposition assay, were a gift from the Boeke laboratory. The following plasmids used for DNA damage experiments in HeLa Tet-On cells were gifts from Kathleen Burns^33^: pDA007 (Addgene plasmid # 131380), pDA025 (Addgene plasmid # 131384), pDA027 (Addgene plasmid # 131385), and pDA034 (Addgene plasmid # 131386).

### Cloning and plasmids

The untagged L1 EN WT protein expression plasmid was generated by restriction digest cloning. The L1 ORF2 consensus sequence was optimized for expression in *E. coli* and synthesized in pUC57 by GenScript. The sequence corresponding to residues 1-239 was amplified by PCR using Q5 High-Fidelity DNA Polymerase (NEB), cut with restriction enzymes NdeI (NEB) and XhoI (NEB), and ligated into digested pET26b with T4 DNA ligase (NEB). Individual colonies were tested by colony PCR and the insert was confirmed by sequencing following Miniprep (Qiagen).

Plasmids expressing *ORFeus* EN domain only were generated by restriction digest cloning. The sequence corresponding to residues 1-239 from pDA007 (WT EN) or from pDA025 (H230A EN) was amplified as described above, cut with BamHI and PacI, and ligated into the digested pDA007 backbone as described above. Inserts were confirmed as described above.

### Cell culture

HeLa Tet-On cells (Takara Bio Inc.) were cultured at 37 °C at 5% CO2 in air atmosphere. The cells were grown in DMEM with 10% FBS, 2mM glutamine, and penicillin and streptomycin. HeLa Tet-On cultures containing pPM404 were maintained in 1ug/mL puromycin to select for plasmid retention. LF1 cells were maintained as previously described^12^. Plasmids pDA007, pDA025, pDA027, pAD034, EN WT, and EN H230A *ORFeus* expression constructs were introduced into HeLa Tet-On cells by transfection with FuGENE HD (Promega) for 24 hours. After removal of FuGENE HD, cells were selected with 1ug/mL puromycin for 2 weeks before freezing. Cultures were tested regularly with MycoAlert Mycoplasma Detection Kit (Lonza).

### Molecular docking and pharmacophore generation

The following programs were used for molecular docking: LeDock^38^, AutoDock Vina^40^, and DOCK 6.9^41^. The published EN WT crystal structure (1VYB) or structure with Mn^2+^ bound was used as indicated. For all programs, mol2 files were obtained from the ZINC database^39^. For AD2 analogs, the compounds were chosen by selecting “Find All” under AD2 “Interesting Analogs” and limiting to “For Sale” compounds. Existing ZINC subsets, such as “fda” and “world-not-fda”, were downloaded directly from ZINC and similarly limited to “For Sale”. The pharmacophore was generated using the EN/DNA co-crystal structure^28^ (PDB ID: 7N94) with ZINCPharmer^42^. Detected features were chosen to reflect key DNA and active site residue interactions and based on similarity to APE-1/DNA pharmacophore features^30^. The final pharmacophore contained 5 features: 2 hydrophobic centers and 3 hydrogen-bond acceptors.

### EN expression, purification, and crystallization

The EN WT plasmid was transformed into BL21 Star (DE3) competent cells (Invitrogen) for large-scale expression. EN WT cultures were grown at 37 °C in 50ug/mL kanamycin until they reached an OD_600_ of 0.6-0.9 and then were induced with 0.5mM IPTG. The cultures were then grown for 2 hours at 37 °C before harvesting by centrifuging at 4000 x g for 12 minutes at 4 °C and then storing at -80 °C.

Cell pellets were resuspended in 10mL EN Lysis Buffer (20mM HEPES, pH 7.5, 300mM NaCl, 1mM DTT) for each 1g of cell pellet and lysed with an Avestin EmulsiFlex C3 (ATA Scientific). Lysate was centrifuged at 30,000 rpm for 1 hour at 4 °C and the supernatant was filtered prior to loading onto a manually packed 20mL Heparin affinity column. Protein was eluted using a gradient of 30%-100% Buffer B (Buffer A: 20mM HEPES, pH 7.5, 1mM DTT; Buffer B: 20mM HEPES, pH 7.5, 1M NaCl, 1mM DTT), diluted to 400mM NaCl, and loaded onto a manually packed Sepharose SP Fast Flow cation exchange column. Protein was eluted using a gradient of 40%-100% Buffer B. Fractions containing protein were pooled, concentrated, filtered, and loaded onto a HiPrep Sephacryl S-100 16/60 size exclusion column. Protein was eluted, aliquoted, and stored at -80 °C in EN Lysis Buffer. Protein purification results were confirmed by SDS-PAGE.

EN WT crystals were grown by mixing an equal volume of 15mg/mL protein in EN Lysis Buffer with an equal volume of crystallizing condition based on the published crystal structure^27^: 0.14M ammonium sulfate, 24% polyethylene glycol (PEG) 5000 monomethyl ether, and 5mM magnesium chloride. Crystals were soaked in cryoprotecting solution containing crystallizing condition, 30% PEG200, and 100mM manganese sulfate before flash freezing in liquid nitrogen. Diffraction images were collected at the NSLS-II AMX beamline at Brookhaven National Laboratory. Images were processed using XDS^56^ and Aimless in CCP4^57^. The published structure^27^ (PDB ID: 1VYB) was used as the search model for molecular replacement with Phaser in Phenix^58^. Anomalous diffraction maps were generated to determine the location of the Mn^2+^ ion. The structure was finished by iterative rounds of manual building in Coot^59^ and refinement in Phenix. Mn^2+^ coordination by active site residues and water molecules was evaluated with CheckMyMetal^60^.

### Plasmid nicking activity assay

The plasmid nicking assay was performed based on a previous assay^47^ with the following modifications. The substrate supercoiled plasmid used was pUC57 containing the *E. coli* codon optimized L1 ORF0 sequence generated by Miniprep (Qiagen). 8nM EN WT and inhibitors or vehicle were incubated at room temperature for 1 hour before adding 2nM plasmid. Reactions were incubated at 37 °C for 3 hours before stopping the reaction with heat inactivation (70 °C for 10 minutes) or addition of 50mM EDTA. Reactions were run on a 1% agarose gel in 1X TAE buffer and visualized with ethidium bromide. Supercoiled plasmid without EN and linearized plasmid were included as controls.

### Fluorescent oligonucleotide nicking activity assay and quantification

The hairpin sequence for the fluorescent oligonucleotide nicking activity assay was adapted from a previous assay^29^. The EN target sequence was added to the stem of the hairpin, the 5-FAM fluorescent tag was included at the 5’ end, and the DABCYL quencher was included at the 3’ end (5’-FAM-CGACTTTTAGATTGACACGCCATGTCGATCAATCTAAAAGTCG-DABCYL-3’). Reactions were completed in buffer containing 20mM HEPES pH 7.5, 50mM NaCl, and 2.5mM MgCl2. EN WT at 2.5nM was incubated with inhibitors or vehicle for 1 hour before adding 25nM oligo. Fluorescence was measured at regular intervals at 37 °C with excitation 485nm and emission 530nm using a Synergy H1 plate reader (BioTek). Initial rates were normalized to no inhibitor control to calculate percent activity and IC_50_ values were obtained using [inhibitor] vs. response non-linear fit in GraphPad Prism version 9.4.1 for Windows. No inhibitor control and full inhibition by 50mM EDTA were included in fit calculations to guide definition of the top and bottom of the fit curve^61^.

### HeLa dual-luciferase L1 retrotransposition assay

The L1 retrotransposition assay was performed using HeLa Tet-On cells containing pPM404. Cells were maintained in 1ug/mL puromycin and seeded into a 96-well plate at a density of 15,000 cells per well. Cells were induced with 1ug/mL doxycycline and treated with inhibitors or vehicle as indicated for 48 hours. Cytotoxicity was evaluated with PrestoBlue Viability Reagent by incubating with cells at 37 °C for 15 minutes and reading fluorescence with excitation 550nm and emission 600nm using a Cytation 5 Plate Reader (BioTek). Luciferase activity was then measured with the Dual-Luciferase Reporter Assay System (Promega) also using the plate reader.

### Immunofluorescence imaging and γ-H2AX quantification

Cells for immunofluorescence were grown on coverslips in 24-well plates and washed with PBS prior to fixation in 4% paraformaldehyde for 20 minutes. Samples were treated with Permeabilization Buffer (PBS, 0.2% Triton) for 20 minutes and treated with Blocking Buffer (PBS, 0.02% Triton, 3% BSA) for 20 minutes. Primary antibody were diluted in Blocking Buffer as described below and incubated with samples for 2 hours: γ-H2AX monoclonal mouse antibody JBW301 (Millipore Sigma, 1:1000). Samples were washed with Blocking Buffer twice for 5 minutes each then treated with secondary antibody diluted 1:200 for 2 hours. Samples were washed twice with PBS for 5 minutes then treated with DAPI (1ug/mL) for 15 to 30 minutes. Finally, coverslips were mounted onto slides with ProLong Gold Antifade Mountant (Invitrogen). Images were acquired with a Nikon Ti2-E Fluorescence Microscope.

HeLa Tet-On cells containing *ORFeus* plasmids as indicated were induced with 2ug/mL doxycycline and treated with inhibitors or vehicle for 24 hours before fixation as described above. Quantification of γ-H2AX was completed with CellProfiler and statistical analysis completed with GraphPad Prism version 9.4.1 for Windows (ROUT outlier correction with Q=0.1% and one-way ANOVA followed by Dunnett’s multiple comparisons test). An existing pipeline was modified to identify nuclei, identify γ-H2AX signal, and measure the mean γ-H2AX intensity for each nucleus with γ-H2AX signal. The pipeline efficacy was confirmed by manual validation of automatic nucleus detection. All images for each independent experiment were acquired during the same imaging session and with the same exposure.

### Neutral COMET assay

HeLa Tet-On cells containing *ORFeus* EN WT or EN H230A plasmids were induced with 2ug/mL doxycycline and treated with inhibitors or vehicle for 24 hours. Neutral COMET assay was performed according to the Trevigen CometAssay Kit instructions with the following modifications. Samples were treated with COMET Assay Lysis Buffer (2.5M NaCl, 100mM EDTA, 10mM Tris, and 0.1% Triton X-100, at pH 10) at 4 °C overnight. Cells were imaged with a Nikon Ti2-E Fluorescence Microscope. All images for each independent experiment were acquired during the same imaging session and with the same exposure.

### Senescent cell culture and RT-qPCR

Replicative senescent LF1 cultures were generated as previously described^12^. Briefly, cultures were passaged until cells reached replicative exhaustion. Cells were designated as entering senescence between 1 week before the date of the last passage, or 1 week after. Media on cells was then replaced twice a week without passaging cells. After 1 month of senescence cells were replated 1:1 into new 10 cm plates in case of contact inhibition. Senescence identity was evaluated with several metrics including the senescence-associated β-galactosidase assay^62^, cell enlargement, RT-qPCR of SASP markers, and immunofluorescence of γ-H2AX and L1 ORF1. Following treatment with inhibitors or no inhibitor control, cultures were harvested for RNA using Trizol (Invitrogen) and cDNA was generated using the TaqMan reverse transcription kit (Applied Biosystems). RT-qPCR was run using the ViiA 7 Real-Time PCR system. Primer sequences used were previously described^12^.

## Acknowledgments

This research was supported by NIH R01 AG016694, NIH P01 AG051449, and NIH 5T32GM007601. Brown University has filed patents on the intellectual property described in this paper.

This research used the AMX beamline of the National Synchrotron Light Source II, a U.S. Department of Energy (DOE) Office of Science User Facility operated for the DOE Office of Science by Brookhaven National Laboratory under Contract No. DE-SC0012704. The Center for BioMolecular Structure (CBMS) is primarily supported by the National Institutes of Health, National Institute of General Medical Sciences (NIGMS) through a Center Core P30 Grant (P30GM133893), and by the DOE Office of Biological and Environmental Research (KP1607011).

The authors would like to thank all members of the laboratories of John Sedivy, Gerwald Jogl, and Jill Kreiling for feedback and support. A.M.D. would specifically like to thank Bianca Kun and Anna Petrashen for assistance with senescent cell culture maintenance. The authors would like to thank Paolo Mita and Jef Boeke for providing the HeLa Tet-On pPM404 cell line. The authors would also like to thank the following core facility managers: Christoph Schorl (Genomic Core Facility), Mandar Naik (Structural Biology and Proteomics Core Facilities), and Geoff Williams (Leduc Bioimaging Facility).

